# Mutation-driven evolution of *Pseudomonas aeruginosa* in the presence of either ceftazidime or ceftazidime/avibactam

**DOI:** 10.1101/359802

**Authors:** Fernando Sanz-García, Sara Hernando-Amado, José Luis Martínez

## Abstract

Ceftazidime/avibactam is a combination of beta-lactam/beta-lactamases inhibitor, which use is restricted to some clinical cases including cystic fibrosis patients infected with multidrug resistant *Pseudomonas aeruginosa*, in which mutation is the main driver of resistance. This study aims to predict the mechanisms of mutation-driven resistance that are selected for when *P. aeruginosa* is challenged with either ceftazidime or ceftazidime/avibactam. For this purpose, *P. aeruginosa* PA14 was submitted to experimental evolution in the absence of antibiotics and in the presence of increasing concentrations of ceftazidime or ceftazidime/avibactam for 30 consecutive days. Final populations were analysed by whole-genome sequencing. All evolved populations reached similar levels of ceftazidime resistance. Besides, all of them were more susceptible to amikacin and produced pyomelanin. A first event in the evolution was the selection of large chromosomal deletions containing *hmgA* (involved in pyomelanin production), *galU* (involved in β-lactams resistance) and *mexXY-oprM* (involved in aminoglycoside resistance). Besides mutations in *mpl* and *dacB* that regulate β-lactamase expression, mutations related to MexAB-OprM overexpression were prevalent. Ceftazidime/avibactam challenge selected mutants in the putative efflux pump *PA14_45890-45910* and in a two-component system *(PA14_45870-45880)*, likely regulating its expression. All populations produce pyomelanin and were more susceptible to aminoglycosides likely due to the selection of large chromosomal deletions. Since pyomelanin-producing mutants, presenting similar deletions are regularly isolated from infections, the potential aminoglycosides hyper-susceptiblity and reduced β-lactams susceptibility of pyomelanin-producing *P. aeruginosa* should be taken into consideration for treating infections by these isolates.

## INTRODUCTION

*Pseudomonas aeruginosa* is an opportunistic pathogen widely distributed in nature (1), which is a major cause of lung and airway infections in hospitalized patients, as well as chronic infections in patients with cystic fibrosis (CF) and chronic obstructive pulmonary disease (2, 3). This bacterial species presents a characteristic low susceptibility to antibiotics, including β-lactams, which is mainly the consequence of its low permeability and the presence in its genome of different intrinsic resistance genes, including those encoding multidrug (MDR) efflux pumps (4) and beta-lactamases. In addition, an increasing number of *P. aeruginosa* isolates has acquired several resistance genes through horizontal gene transfer (HGT), including different classes of carbapenemases. Finally, *P. aeruginosa* is able to develop resistance through mutation, particularly when causing chronic infections, to nearly any available antibiotic. Under this situation, the emergence and spread of MDR resistant global clones is of special concern (5).

The use of β-lactam/β-lactamase inhibitor combinations, such as amoxicillin/clavulanic acid or ceftolozane/tazobactam, has proven to be effective against class A β-lactamases (which include narrow and extended-spectrum β-lactamases and some carbapenemases); whereas effective combinations against class B, C (extended spectrum cephalosporinases) and D β-lactamases (6-8) have not been available until recently. One of them is the ceftazidime/avibactam combination, whose use was approved in 2015 by the FDA (9).

Avibactam, formerly known as NXL104, belongs to a new class of β-lactamase inhibitors, the diazabicyclooctanes (10). This inhibitor has a potent activity against most Class A, Class C and some Class D β-lactamases (11). Avibactam has been mainly used for restoring the activity of the third generation cephalosporin ceftazidime (12). Thus far, it has been used for the treatment of patients with complicated urinary tract infections, including pyelonephritis; and community-acquired intra-abdominal infections, usually in combination with metronidazole (13). Besides, future studies are likely to expand the use of ceftazidime/avibactam to include other cases, such as cystic fibrosis patients with MDR resistant *P. aeruginosa* infections (13).

Given the fact that this treatment is currently reserved for patients who have no alternative therapeutic options, a judicious use of antibiotic stewardship should be applied in order to prevent the incidence of drug resistance. Nevertheless, and although there are numerous studies on the activity of ceftazidime/avibactam against pathogens resistant to other antibiotics (14-17), analysis for predicting potential mechanisms of resistance to this antimicrobial combination are still scarce.

In the present study, experimental evolution followed by whole-genome sequencing (WGS) was used to examine the evolutionary trajectories taken by *P. aeruginosa* towards resistance against the combination ceftazidime/avibactam compared to the ones followed in absence of avibactam. This may throw light upon the different mechanisms of resistance that are selected for in *P. aeruginosa* when its β-lactamase activity is inhibited by the presence of this novel inhibitor. In addition, the present work may allow us to elucidate whether the presence of avibactam modifies the resistance level acquired by the bacterial populations in comparison to the one developed when ceftazidime is used alone. Thus, these results may give rise to strategies for predicting, managing, and eventually reducing resistance to ceftazidime/avibactam. This is widely important, as this treatment is strictly restricted to few clinical cases in which resistant strains would be of major concern.

## MATERIALS AND METHODS

### Growth conditions and antibiotic susceptibility assays

Unless otherwise stated, bacteria were grown in Luria Bertani (LB) Broth at 37°C with shaking at 250 rpm. The susceptibility to tigecycline, tetracycline, aztreonam, ceftazidime, imipenem, meropenem, ciprofloxacin, levofloxacin, norfloxacin, tobramycin, streptomycin, amikacin, gentamycin, colistin, polymyxin B, chloramphenicol, fosfomycin and erythromycin was determined by disk diffusion in Mueller Hinton Agar (MHA) (Sigma) at 37°C. For a set of antibiotics, MICs were determined using E-test strips (MIC Test Strip, Liofilchem®). MICs of ceftazidime and ceftazidime/avibactam were determined in LB by double dilution in microtiter plates.

### Experimental evolution procedure

Twelve bacterial populations from a stock *P. aeruginosa* PA14 culture (four controls without antibiotic, four populations challenged with ceftazidime, and four populations challenged with ceftazidime/avibactam) were grown in parallel in LB for 30 consecutive days. Each day, the cultures were diluted (1/250) in fresh LB. The concentrations of ceftazidime used for selection were increased over the evolution experiment from the concentration that hinders the growth of *P. aeruginosa* PA14 under these culture conditions (4 μg/ml) up to 128 μg/ml, doubling them every 5 days. The avibactam concentration was maintained constant, as used in clinical tests (18), at 4 μg/ml. In some occasions, the cultures did not grow when antibiotic concentration increased, in which case, the selection was kept at the concentration that allows growth. Every five days, samples from each culture were preserved at -80 °C for further research.

### Whole-genome sequencing (WGS)

Gnome^®^ DNA kit (MP Biomedicals) was used to extract genomic DNA. WGS was performed by Sistemas Genómicos S.L. The quality of the extracted material was analysed via a 4200 Tape Station, High Sensitivity assay and the DNA concentration ascertained by real-time PCR using a LightCycler 480 device (Roche). Libraries were obtained without amplification following Illumina protocols and were pair-end sequenced (100 x 2) in an Illumina HiSeq 2500 sequencer. The average number of reads per sample was 7,178,870, which represents a 200x coverage, on average.

### Bioinformatics analysis of WGS and confirmation of genetic changes

Mutations in the evolved populations were identified using CLC Genomics Workbench 9.0 (QIAGEN). *P. aeruginosa* UCBPP-PA14 reference chromosome (NC_008463.1) was used to align the reads obtained from WGS data (previously trimmed). Sanger sequencing was used to verify and to settle the order of appearance of the putative mutations found via WGS (Table S1). Thirty-two pairs of primers, which amplified 200-400 base pair regions containing each genetic modification, were designed (Table S2). After PCR amplification, the corresponding amplicons were purified using the QIAquick PCR Purification Kit (QIAGEN) and sequenced at GATC Biotech.

## RESULTS

### Stepwise evolution of *P.* aeruginosa towards ceftazidime and ceftazidime/avibactam resistance

To determine the potential evolutionary trajectories that can lead to either ceftazidime or ceftazidime/avibactam resistance, four biological replicates were allowed to evolve in parallel in each of the following conditions (Figure 1): under selective pressure with ceftazidime (populations 1-4), ceftazidime/avibactam (populations 5-8), and in the absence of any selective pressure (populations 9-12). The susceptibility of each population to the selecting antibiotic was determined every 5 days by E-test. However, after 20 days of evolution, MICs reached the highest limits of the E-test strips and MICs were again determined, for each evolutionary step, by double dilution (Table S3). Stepwise evolutionary trajectories were observed for both treatments, in which the selected populations reached quite similar levels of resistance (Figure 2). These results suggest that avibactam inhibition may not be a guarantee of impeding *P. aeruginosa* to acquire high-level ceftazidime resistance. An increase in the MIC of an antibiotic after experimental evolution does not necessarily imply that antibiotic-resistant mutants have been selected for: resistance may have arisen due to a phenotypic (inducible) adaptation to the presence of ceftazidime rather than to mutations (19-21). To address this possibility, the evolved populations were cultured in the absence of selection pressure (three sequential passages on LB) and the MICs again determined. These were found not to vary, indicating that the observed modifications were due to the selection of stable mutants.

**Figure 1.**
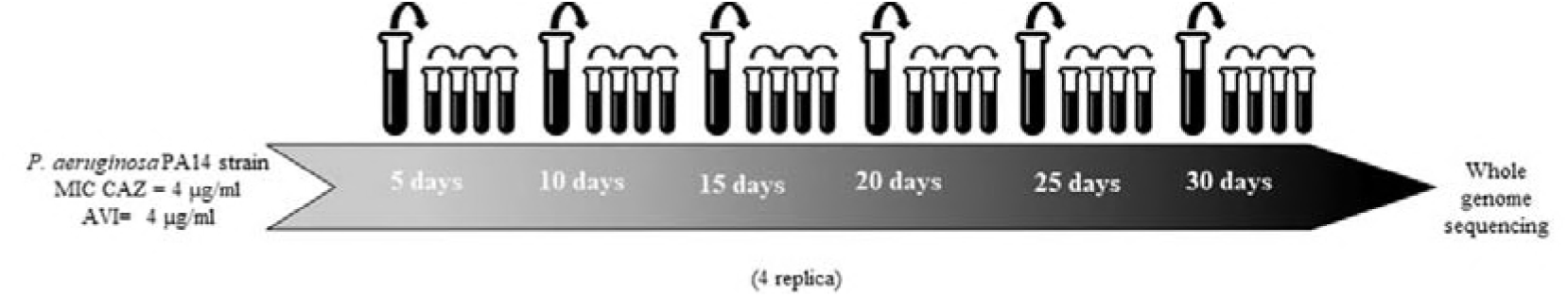
Experimental evolution assay. Eight bacterial cultures of *P. aeruginosa* PA14 strain were challenged with increasing inhibitory concentrations of ceftazidime in Luria Bertani Broth (LBB) for 30 consecutive days. The ceftazidime concentration was raised by two-fold every five days, from 4 μg/ml up to 32MIC. Avibactam was added in combination with ceftazidime in four of these eight populations, at constant concentration of 4 μg/ml, as it is the value established in clinical tests. Four controls without any selective pressure were also grown in parallel. At the end of the experimental evolution, the genomic DNA of the twelve independent populations was extracted and analysed by whole-genome sequencing (WGS).

**Figure 2.**
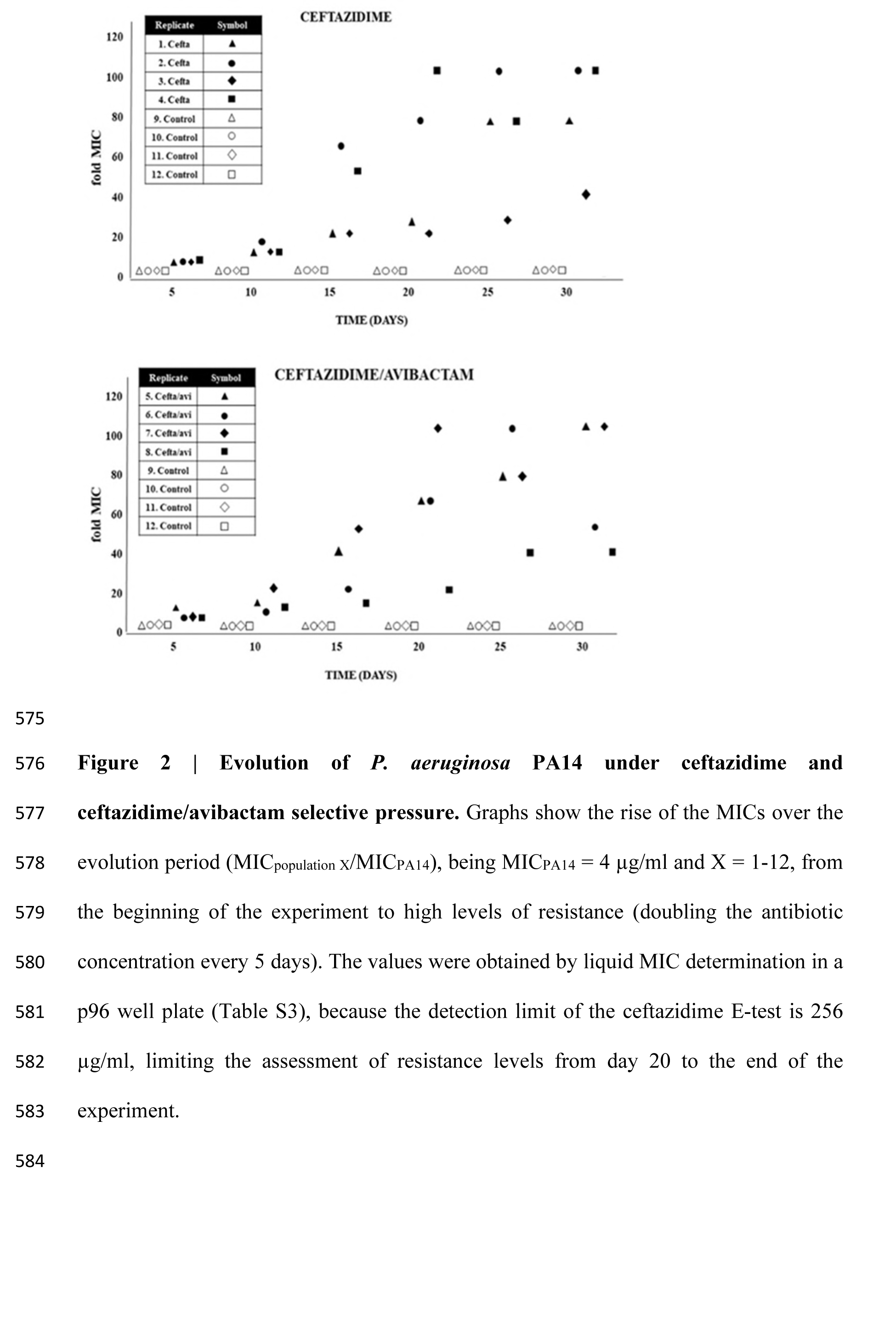
Evolution of *P. aeruginosa* PA14 under ceftazidime and ceftazidime/avibactam selective pressure. Graphs show the rise of the MICs over the evolution period (MIC_population_X/MICPA14), being MIC_PA14_ = 4 μg/ml and X = 1-12, from the beginning of the experiment to high levels of resistance (doubling the antibiotic concentration every 5 days). The values were obtained by liquid MIC determination in a p96 well plate (Table S3), because the detection limit of the ceftazidime E-test is 256 μg/ml, limiting the assessment of resistance levels from day 20 to the end of the experiment.

### Cross-resistance and collateral sensitivity of the evolved populations

Taking into consideration the few therapeutic options for patients submitted to ceftazidime/avibactam therapy, knowing whether or not acquisition of resistance to this combination might alter the susceptibility to other antibiotics is of crucial importance. To that end, the susceptibility to a range of representative antibiotics was tested by disk diffusion assay (Table S4). From these results, a set of antibiotics was chosen for determining their MICs against the different evolved populations. Every evolved replicate showed altered susceptibility to antimicrobials belonging to different structural families (Table 1), implying that at least some resistance mutations were not ceftazidime- or ceftazidime/avibactam-specific. All populations evolved in the presence of either ceftazidime or ceftazidime/avibactam presented decreased susceptibility to other β-lactams, to chloramphenicol and to erythromycin and they were more susceptible to fosfomycin and to amikacin. Notably, while populations evolving in the presence of ceftazidime were less susceptible to tetracycline and did not present changes in the susceptibility to tigecycline, populations evolved in the presence of ceftazidime/avibactam were hyper-susceptible to both antibiotics.

**Table 1.** MICs (µg/ml) of antibiotics of different structural families in the ι populations evolved in the presence of either cefatzidime or ceftazidime/avibactam. Populations challenged with ceftazidime: 1-4, ceftazidime/avibactam: 5-8, and controls: 9-12. Tgc: tigecycline, tet: tetracycline, atm: aztreonam, caz: ceftazidime, cza: ceftazidime+avibactam, ipm: imipenem, mer: meropenem, nor: norfloxacin, ak: amikacin, c: chloramphenicol, e: erythromycin, f: fosfomycin.

| Replicate | Tgc | Tet | Atm | Ipm | Mer | Caz | Cza | Nor | Ak | C | E | F |
| --- | --- | --- | --- | --- | --- | --- | --- | --- | --- | --- | --- | --- |
| **PA14** | 2 | 12 | 1.5 | 0.5 | 0.19 | 1 | 1 | 0.25 | 1.5 | 24 | 32 | 16 |
| **1** | 2 | 32 | ≥256 | 8 | 32 | ≥256 | ≥256 | 1.5 | 1 | ≥256 | ≥256 | 4 |
| **2** | 1.5 | 24 | ≥256 | 4 | 32 | ≥256 | ≥256 | 1 | 0.5 | ≥256 | ≥256 | 1.5 |
| **3** | 2 | 16 | ≥256 | 2 | 32 | ≥256 | ≥256 | 1 | 0.75 | ≥256 | ≥256 | 4 |
| **4** | 2 | 16 | ≥256 | 3 | 32 | ≥256 | ≥256 | 1 | 0.5 | ≥256 | ≥256 | 2 |
| **5** | 0.75 | 4 | ≥256 | 32 | 32 | ≥256 | ≥256 | 0.25 | 0.75 | ≥256 | ≥256 | 3 |
| **6** | 0.5 | 4 | ≥256 | 3 | 16 | ≥256 | ≥256 | 0.25 | 1 | ≥256 | ≥256 | 4 |
| **7** | 0.38 | 4 | 96 | 32 | 32 | ≥256 | ≥256 | 0.25 | 2 | ≥256 | ≥256 | 6 |
| **8** | 0.75 | 8 | ≥256 | 4 | 32 | ≥256 | ≥256 | 0.75 | 0.75 | ≥256 | ≥256 | 4 |
| **9** | 3 | 12 | 1.5 | 0.75 | 0.25 | 1 | 1 | 0.25 | 1.5 | 24 | 32 | 24 |
| **10** | 3 | 12 | 1.5 | 0.5 | 0.19 | 1 | 1 | 0.25 | 1.5 | 24 | 32 | 24 |
| **11** | 3 | 12 | 1 | 0.75 | 0.25 | 1 | 1 | 0.25 | 1.5 | 24 | 32 | 16 |
| **12** | 3 | 16 | 1 | 0.75 | 0.25 | 1 | 1 | 0.38 | 1.5 | 24 | 32 | 12 |

### Analysis of mutations associated with the acquisition of resistance

To know the genetic events associated with the acquisition of resistance in the evolved populations, the genomes of each, as well as that of the original PA14 strain, were sequenced on the last day of the experiment. Table 2 encompasses the resulting mutated genes and their functional significance, whereas Table S1 shows the locations of all 40 genetic changes that were unveiled and were not present in control populations evolving in the absence of antibiotics. A total of 37 single nucleotide variants (SNVs) and 3 multi-nucleotide variants (MNVs; deletions and substitutions of various nucleotides) were found, 36 located in genes and 4 in intergenic regions. Most mutations located in genes resulted in amino acid alterations, stop codons or frameshifts. In addition, all the populations evolved in the presence of antibiotics contained large chromosomal deletions (55 to 443 kbp) representing from 0,88% to 7,09% of the *P. aeruginosa* PA14 genome. Five different deletions were selected, they all presenting a 55 kbp common region (Figure 3).

**Figure 3.**
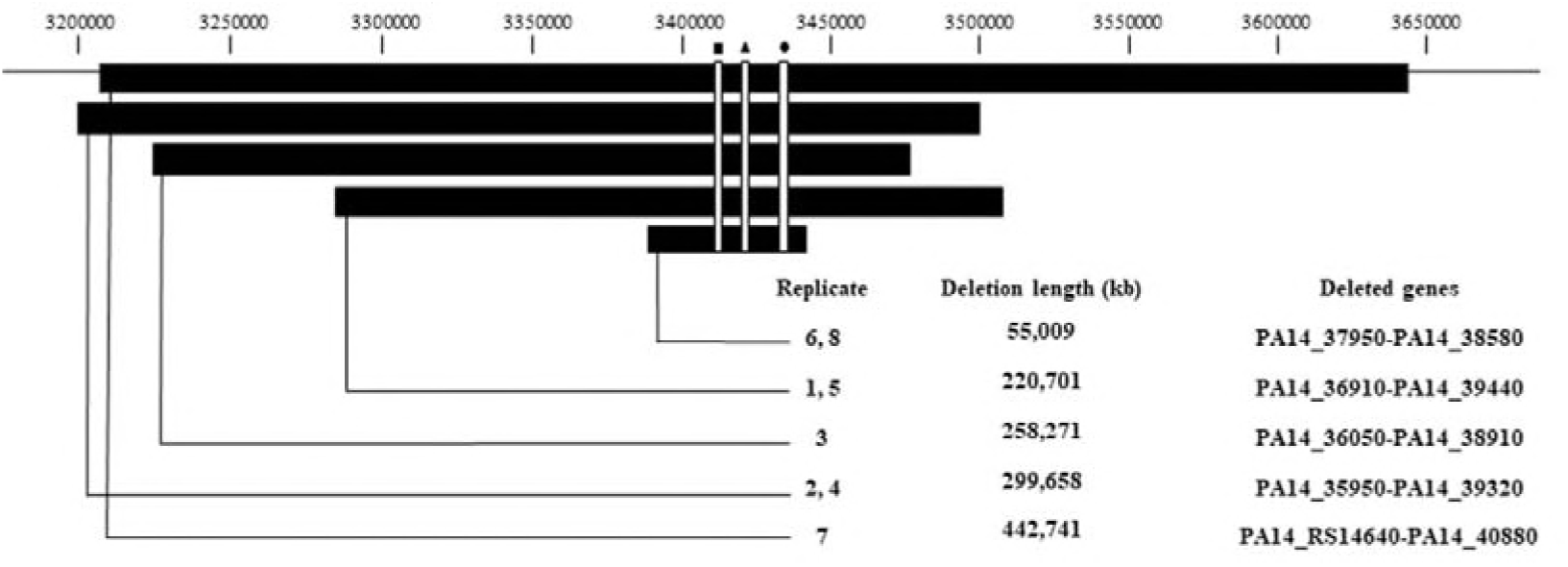
Large deletions present in all ceftazidime and ceftazidime/avibactam evolved populations since the first day of the experimental evolution. The figure indicates the length of the deletions and the deleted genes in each replicate, as well as their genome localization, which corresponds with *P. aeruginosa* UCBPP-PA14 reference chromosome (NC_008463.1). Black square: *mexXY*, black triangle: *galU*, black circle: *hmgA.*

**Table 2.**
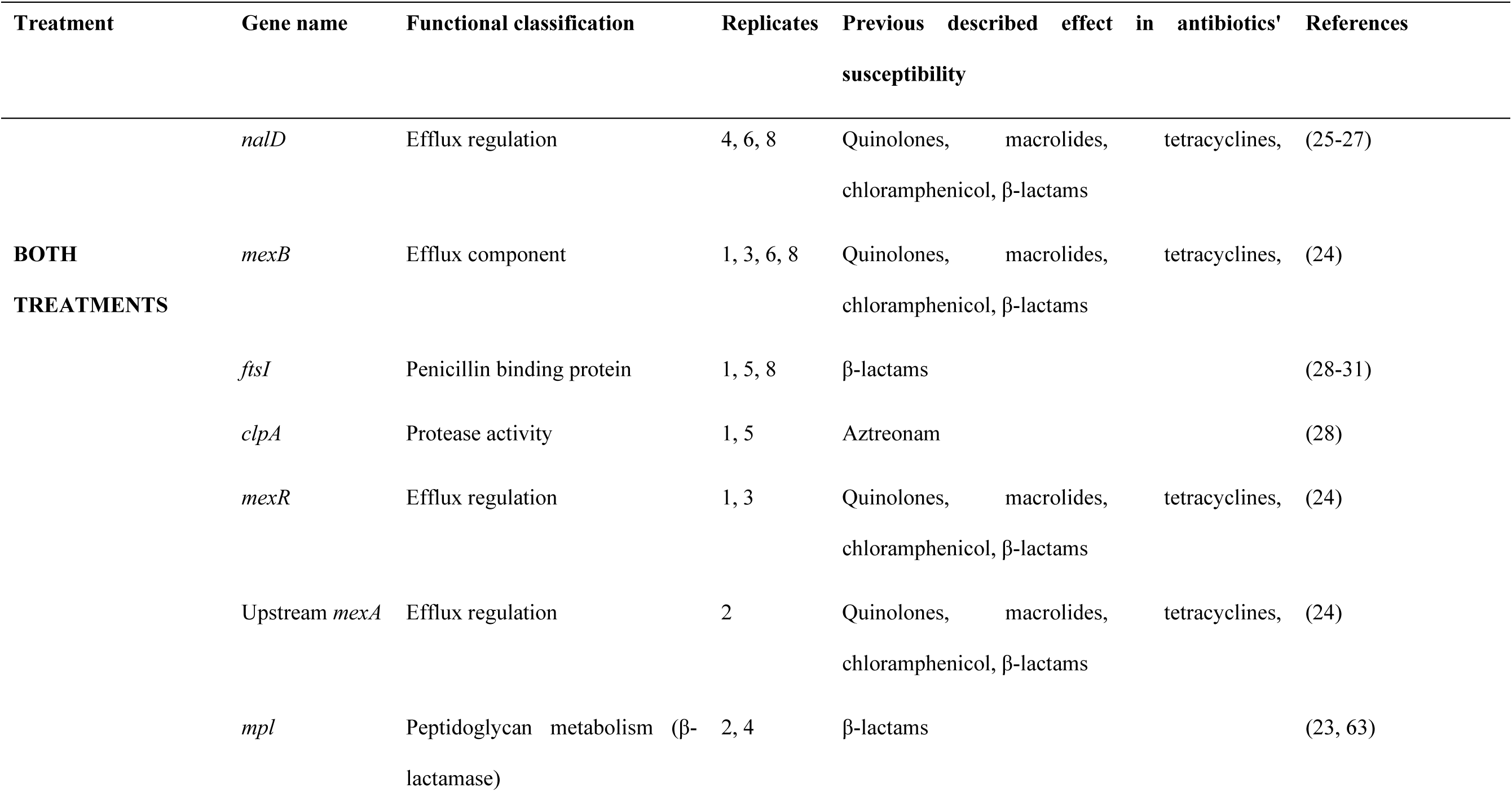
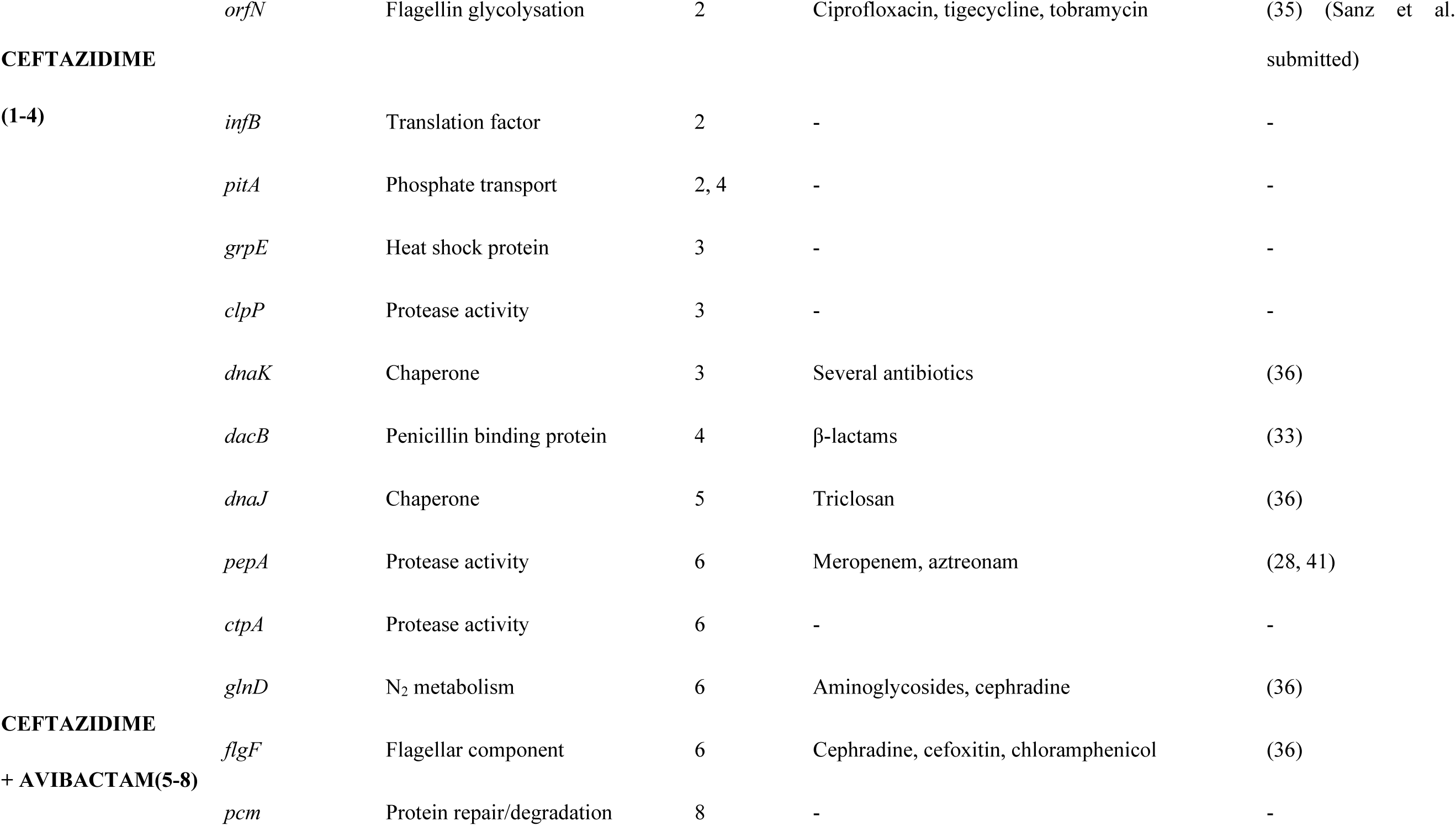
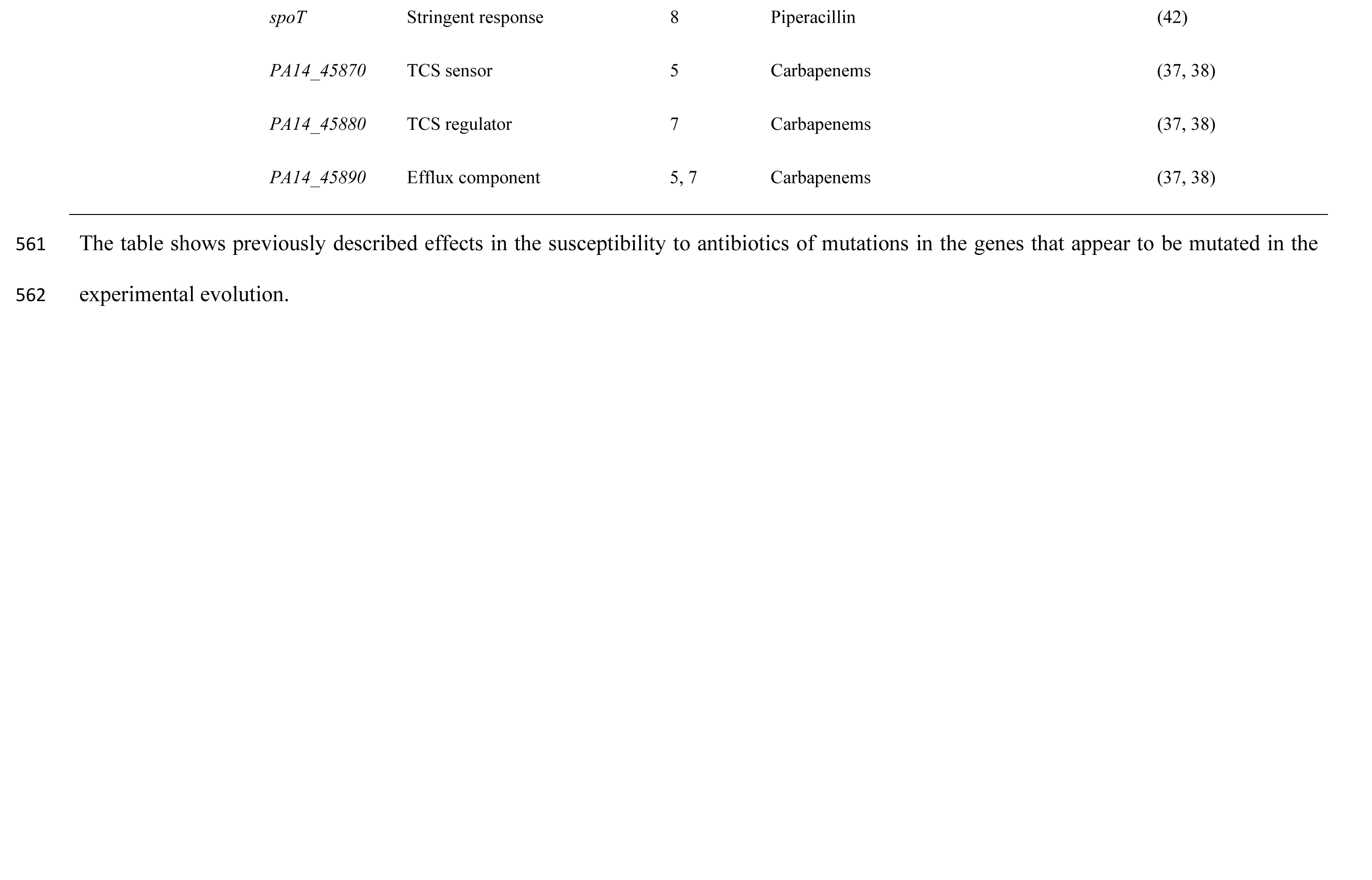
Mutated genes in ceftazidime and ceftazidime/avibactam evolved populations. The table shows previously described effects in the susceptibility to antibiotics of mutations in the genes that appear to be mutated in the experimental evolution.

To verify the presence and the order of appearance of the genetic changes identified by WGS, the regions holding these mutations were amplified using specific oligonucleotides (Table S2) and the amplicons Sanger-sequenced in each evolutionary step (Figure 4). Regarding the large chromosomal deletions, primers located at the flanking sequences were used to verify their presence. In all cases, these analyses confirmed the information obtained from WGS.

**Figure 4.**
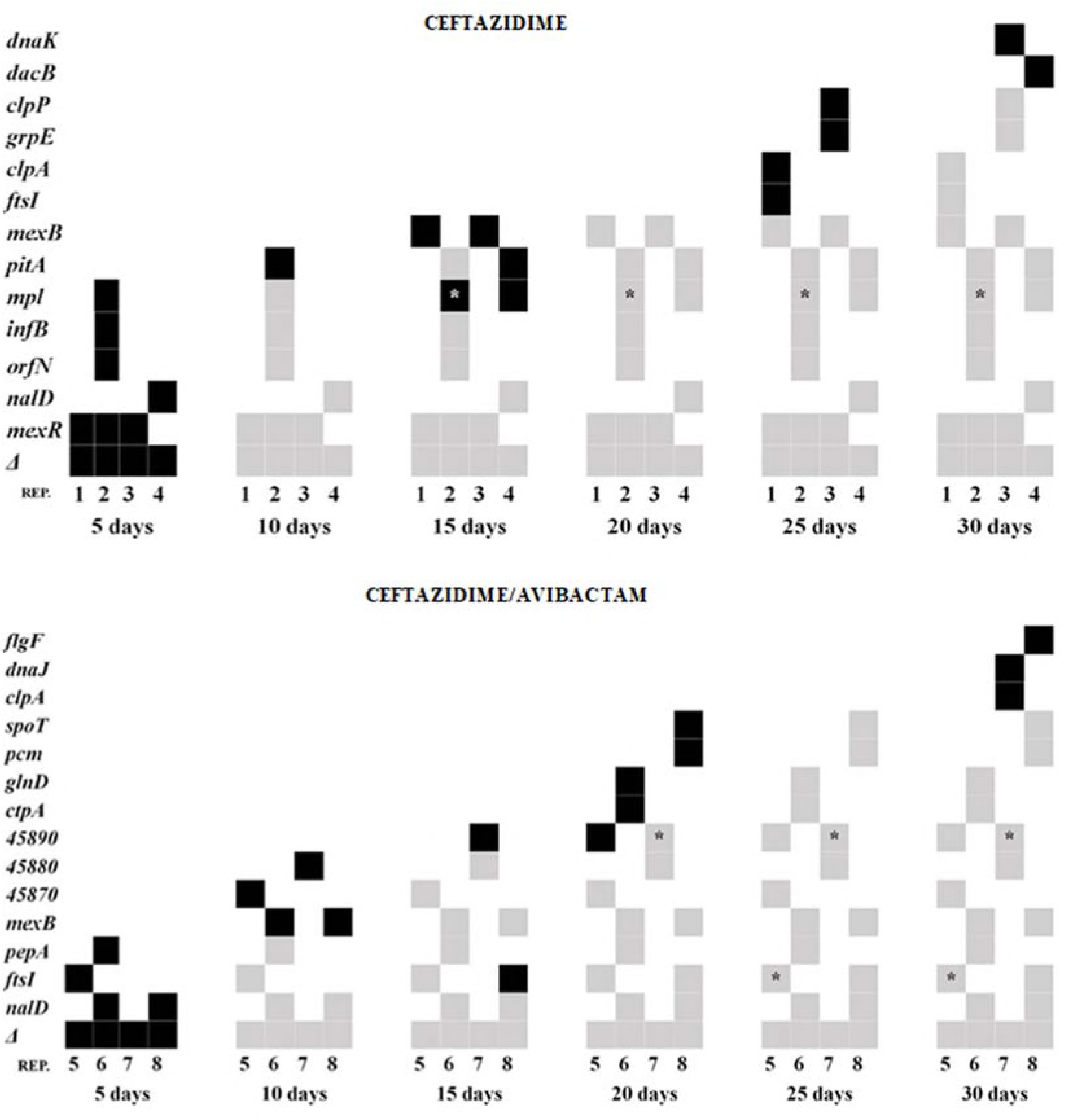
Order of appearance of genetic changes. Ceftazidime and ceftazidime/avibactam resistance mutations appearance during the evolution process, as determined by PCR amplifications of known SNVs/MNVs in evolved populations. The *mexR* mutation in ceftazidime population 2 actually indicates that this mutation occurred in the intergenic region between *mexR* and *mexA.* The “*” refers to a second mutation in a gene that mutated before. We cannot ditch that other mutations may have appeared over the 30 days evolution process. “Δ”: large deletion. Rep: replicate. Black square: the mutation appeared in this evolutionary step. Grey square: the mutation appeared in a previous evolutionary step.

### Common mutations selected upon either ceftazidime or ceftazidime/avibactam selection pressure

Upon one day of experimental evolution, all *P. aeruginosa* PA14 cultures challenged with antibiotic produced a brown pigment, which appeared to be pyomelanin, whose accumulation is normally due to the lack of homogentisate 1,2-dioxygenase activity provided by the enzyme HmgA (22). All the chromosomal large deletions selected during evolution presented *hmgA* (Figure 3). In addition, the deletions included as well *galU* (involved in LPS biosynthesis), whose inactivation reduces ceftazidime susceptibility (23); and the MDR efflux pump *mexXY-oprM*, which contributes to aminoglycoside resistance in *P. aeruginosa* (24). Deletion of the latter is likely the cause of the observed amikacin hyper-susceptibility of all evolved populations (Table 1).

Another common element in both evolutions is *nalD*, which encodes a secondary repressor of MexAB-OprM (25, 26). Three out of eight replicates showed the same T11N change, which has been previously found in XDR *P. aeruginosa* high-risk clones (27) that overexpress MexAB-OprM. Notably, four replicates (including the three presenting mutations in *nalD)* also presented mutations in *mexB*, indicating this efflux system to be a relevant element in the acquisition of resistance. Two other elements that are selected for in both treatments are *ftsI* and *clpA.* The first encodes PBP3, the target of different β-lactams (28, 29), which has been already found to be mutated in numerous resistant *P. aeruginosa* isolates. Indeed, the mutations R504C/H found in populations 1 and 5 are also present among isolates from widespread nosocomial P. *aeruginosa* clones (29-31). *clpA* encodes for an intracellular protease involved in different aspects of *P. aeruginosa* physiology, in addition to aztreonam resistance (28, 32).

### Mutations selected by ceftazidime

In addition to the observed mutations in *nalD*, which would allow *mexAB-oprM* overexpression, we found as well mutations that should lead to the overexpression of this system in the populations evolving under ceftazidime challenge. Two populations carried mutants in *mexR*, which encodes a local repressor of *mexAB-oprM* expression. Another population presented a mutation upstream *mexA* that might prevent the interaction of NalD with its operator (25), thus allowing *mexAB-oprM* overexpression.

Other mutations specifically selected for by ceftazidime were found in *mpl* and *dacB.* The proteins encoded by these genes are involved in the recycling of peptidoglycan muropeptides. Besides, they control the activity of AmpR and consequently the level of AmpC expression (23, 33), which is known to be a main element in *P. aeruginosa* resistance to β-lactams. Interestingly, *mpl* V124G (replicate 2, Table S1) has been found before in a clinical isolate *(P. aeruginosa* NCGM1984). These finding, along with the *ftsI* and *nalD* mutations aforementioned, validate our experimental evolution approach as a valuable predictive model for the *in vivo* selection of antibiotic resistance.

Finally, mutations at *orfN, pitA, infB, grpE, clpP* and *dnaK* were selected for in populations challenged with ceftazidime. *orfN* codes for a putative glycosyl transferase of type A flagellins (34). Mutations on this gene have been found in ciprofloxacin *P. aeruginosa* resistant strains (35), and also in *P. aeruginosa* populations submitted to tigecycline and tobramycin experimental evolutions (Sanz-García *et al.* submitted). *pitA* encodes a phosphate transporter, *infB* the translation initiation factor IF-2 and *dnaK, grpE* and *clpP* encode proteins involved in regulatory gene networks involved in response to stress. None of them has been previously related to ceftazidime resistance, excepting *dnaK*, whose inactivation leads to stronger susceptibility to various antimicrobials in *Escherichia coli* (36).

### Mutations selected by ceftazidime/avibactam

The challenge with ceftazidime/avibactam selected mutants in a predicted efflux pump *(PA14_45890-45910)*, as well as in the two-components system (TCS) encoded by the operon *PA14_45870-45880*, likely regulating its expression. Previous studies have shown this efflux pump to be involved in *P. aeruginosa* intrinsic resistance (37) and susceptibility to carbapenems (38). Regarding the substrate recognition profile this pump might display, it is remarkable that populations 5 and 7, which present the aforementioned mutations, show a much lower susceptibility to imipenem than any other replicate (Table 1), suggesting this pump to have certain specificity to carbapenems.

Other mutations that were selected upon ceftazidime/avibactam treatment were found in *pepA, spoT, dnaJ* and *flgF. pepA* encodes for a protease necessary for *P. aeruginosa* cytotoxicity, virulence and, consequently, lung infection (39, 40). Although its implication in antibiotic resistance has not been studied in detail, it has been reported that its inactivation confers meropenem resistance in *P. aeruginosa* (41). Moreover, *pepA* mutants are selected in the presence of aztreonam (28). SpoT has been related to piperacillin resistance (42); while DnaJ, a chaperone protein and FlgF, a flagellar basal body rod protein (43), have been reported to modify the susceptibility of *E. coli* to a range of antibiotics when they are inactivated (36).

The other mutations that were selected for in populations under ceftazidime/avibactam challenge; namely those occurring in *ctpA*, an essential gene for the transition between acute and chronic *P. aeruginosa* infection (44), *pcm*, that encodes for a L-isoaspartate carboxylmethyltransferase type II that participates in protein repair and degradation and *glnD*, which is implicated in N2 metabolism, (45) have not been reported to be involved in antibiotic resistance.

## Discussion

The use of β-lactamase inhibitors has re-emerged as a fruitful strategy for fighting infections by MDR bacteria. Among them, ceftazidime/avibactam can be a useful combination for treating infections by different organisms, including *P. aeruginosa.* The analysis of the mechanisms of resistance to previous β-lactam/β-lactamase inhibitor combinations as amoxicillin/clavulanate have shown that the main mechanisms selected along their use have been increased expression or mutation of pre-existing β-lactamases and acquisition of new ones by HGT (46-50). *P. aeruginosa* has already acquired different carbapenemases that might be important elements in ceftazidime/avibactam resistance. In addition, resistance can be achieved through mutations, particularly in the case of *P. aeruginosa* causing chronic infections. To identify potential mutations involved in the acquisition of either ceftazidime or ceftazidime/avibactam resistance, bacterial populations were submitted to increasing selective concentrations of these antimicrobials. In both cases, the first event in the evolution seems to be the deletion of large regions of *P. aeruginosa* chromosome that comprise, among several other genes, *hmgA, galU* and *mexXY.* A similar situation has been previously reported in other *P. aeruginosa* experimental evolution assays in the presence of β-lactams, such as piperacillin (42) and meropenem (51). Additionally, pyomelanin-producing mutants are regularly isolated from infections; up to 13% of CF patients harbour pyomelanin-producing mutants (52), likely because the production of pyomelanin increases resistance to oxidative stress and persistence in chronic lung infections (22). Recent work has also shown that these mutations can be selected to prevent bacteriophage predation (53). Notably, melanogenic clinical isolates of *P. aeruginosa* present large chromosomal deletions, similar to those reported in the present work (54). Our results then support that ceftazidime selects for these genome deletions, and the presence of avibactam cannot prevent them from happening. It might be possible that deletions are the consequence of increased recombination triggered by the presence of the antibiotic. However, the fact that *P. aeruginosa* evolving in the presence of ciprofloxacin do not produce pyomelanin (a marker of these deletions) and that pyomelanin-producing mutants are selected when a *recA P. aeruginosa* defective strain is challenged with either ceftazidime or ceftazidime/avibactam (data not shown), goes against this possibility. Besides the already known effect of the lack of *galU* on the susceptibility to β-lactams, the absence of other genes located in the deletion, such as *mexXY*, may affect *P. aeruginosa* susceptibility to antibiotics. Deletion of this pump is likely the cause of the observed hyper-susceptibility to amikacin of the evolved populations. In addition, it might have an indirect effect on the decreased susceptibility to beta-lactams, particularly in the case of those strains carrying mutations in the repressors of *mexAB-oprM.*

MexAB-OprM is an important determinant of intrinsic *P. aeruginosa* resistance to different antibiotics, including β-lactams (24). Further, mutants overexpressing this efflux pump are regularly isolated from infections and it has been shown its expression to be prevalent among resistant *P. aeruginosa* clinical isolates (55-58). MexAB and MexXY share the outer membrane protein OprM, which produces antagonistic interactions when both systems are expressed (51, 59). Hence, MexXY-OprM elimination might favour the efficiency of β-lactams efflux, reducing the competition of both efflux pumps for OprM.

Important elements in the acquisition of ceftazidime resistance are efflux pumps, particularly MexAB-OprM, since mutations either in the elements regulating its expression or in the efflux pump itself were found in six out of eight evolved populations, whereas the two remaining populations harboured mutants in the putative *PA14_45890-45910* and in its potential TCS regulator. While the substrates of MexAB-OprM are known and include β-lactams, the substrates of *PA14_4590-45910* are unknown. Nevertheless, it is remarkable that populations presenting mutations on this determinant display a much lower susceptibility to imipenem than any other replicate, suggesting this pump to have certain specificity to β-lactams.

Mutations in elements involved in the regulation of AmpC expression were selected when just ceftazidime was used for selection and not in the presence of ceftazidime/avibactam. This suggests that, at least in the *P. aeruginosa* PA14 background, the efficient inhibition by avibactam of intrinsic β-lactamases, preclude the emergence of mechanisms based on their overexpression and other mechanisms, including the above mentioned large deletions and modifications in the activity of efflux pumps are preferentially selected. This does not necessarily mean that resistance to ceftazidime/avibactam cannot be associated to changes in the activity of AmpC, particularly if the challenged isolate is already resistant to ceftazidime. Indeed, avibactam resistant mutants presenting changes in the avibactam binding pocket of AmpC are selected *in vitro* at low frequency from Amp C-overexpressing ceftazidime resistant *P. aeruginosa* isolates (60).

Although most of the mutants here reported have been previously associated to be involved in antibiotic resistance, it is still possible that some of the mutations might be selected for compensating the fitness costs associated with the acquisition of resistance. This might be the case of *ctpA, pcm* or the mutations at structural elements of efflux pumps that were selected after mutations in the regulators of their expression. For the latter, it might also be possible that these mutations increase the capability of extruding the antibiotic substrates, as described for AcrB (61). The fact that in all evolved populations mutants in efflux pumps are selected, provides an explanation of the cross-resistance phenotype observed in all resistant strains. This situation might be of concern, since both ceftazidime and ceftazidime/avibactam might select for resistance to other antibiotics, at least along chronic infections in which mutation is the main cause of acquisition of resistance.

*P. aeruginosa* evolution in chronic infections frequently involves large genome deletions (62), usually linked to the production of pyomelanin (54). Whether these deletions are selected by antibiotic treatment or are just the consequence of the adaptation to the environment of the lungs of the CF patient remains to be established. However, this evolution provides a link between antibiotic resistance and virulence for this relevant pathogen. In any case, and given that deletions containing *galU* and *hmgA* appear to be a first step on the evolution towards ceftazidime/avibactam resistance, pyomelanin production could be considered as a marker in the selection of the antibiotic of choice for treating *P. aeruginosa* infections. Both *in vitro* work, including the results here shown, and the analysis of clinical pyomelanin-producers, have shown that these isolates are hyper-susceptible to aminoglycosides probably because the deletions they present include *mexXY.* It would then be judicious using aminoglycosides, and not β-lactams for treating infections by pyomelanin-producing *P. aeruginosa.*

## Acknowledgments

Work in our laboratory is supported by grants from the Instituto de Salud Carlos III (Spanish Network for Research on Infectious Diseases [RD16/0016/0011]), from the Spanish Ministry of Economy and Competitivity (BIO2017-83128-R) and from the Autonomous Community of Madrid (B2017/BMD-3691). The funders had no role in study design, data collection and interpretation, or the decision to submit the work for publication. FSG is the recipient of a FPU fellowship.

---

**Supplementary Table S1.**
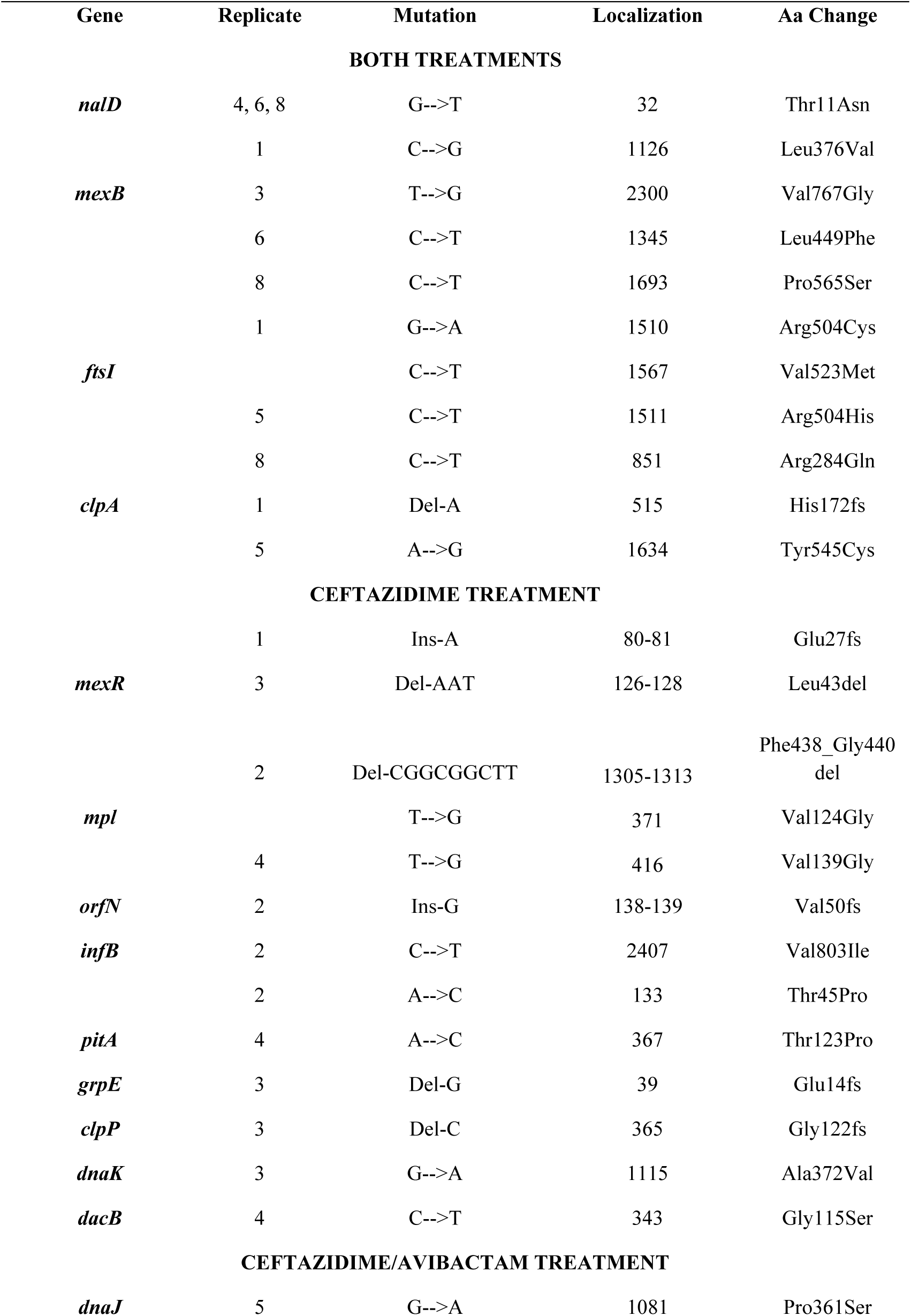
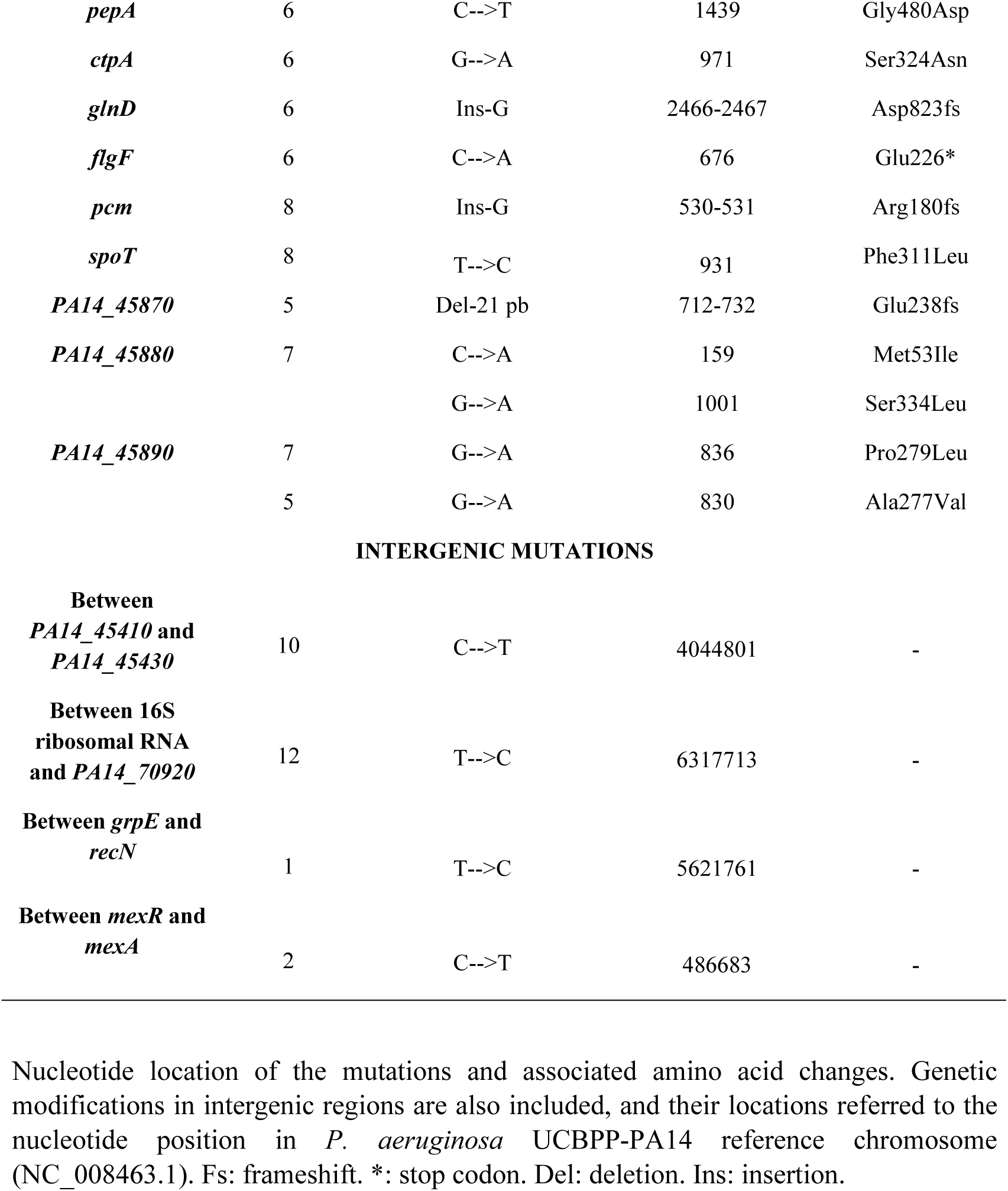
Genetic modifications detected in ceftazidime and ceftazidime/avibactam evolved *P. aeruginosa* PA14 populations. Nucleotide location of the mutations and associated amino acid changes. Genetic modifications in intergenic regions are also included, and their locations referred to the nucleotide position in *P. aeruginosa* UCBPP-PA14 reference chromosome (NC_008463.1). Fs: frameshift. *: stop codon. Del: deletion. Ins: insertion.

| Gene | Replicate | Mutation | Localization | Aa Change |
| --- | --- | --- | --- | --- |
| **BOTH TREATMENTS** |  |  |  |  |
| *nalD* | 4, 6, 8 | G-->T | 32 | Thr11Asn |
|  | 1 | C-->G | 1126 | Leu376Val |
| *mexB* | 3 | T-->G | 2300 | Val767Gly |
|  | 6 | C-->T | 1345 | Leu449Phe |
|  | 8 | C-->T | 1693 | Pro565Ser |
|  | 1 | G-->A | 1510 | Arg504Cys |
| *ftsI* |  | C-->T | 1567 | Val523Met |
|  | 5 | C-->T | 1511 | Arg504His |
|  | 8 | C-->T | 851 | Arg284Gln |
| *clpA* | 1 | Del-A | 515 | His172fs |
|  | 5 | A-->G | 1634 | Tyr545Cys |
| **CEFTAZIDIME TREATMENT** |  |  |  |  |
|  | 1 | Ins-A | 80-81 | Glu27fs |
| *mexR* | 3 | Del-AAT | 126-128 | Leu43del |
|  | 2 | Del-CGGCGGCTT | 1305-1313 | Phe438_Gly440 del |
| *mpl* |  | T-->G | 371 | Val124Gly |
|  | 4 | T-->G | 416 | Val139Gly |
| *orfN* | 2 | Ins-G | 138-139 | Val50fs |
| *infB* | 2 | C-->T | 2407 | Val803Ile |
|  | 2 | A-->C | 133 | Thr45Pro |
| *pitA* | 4 | A-->C | 367 | Thr123Pro |
| *grpE* | 3 | Del-G | 39 | Glu14fs |
| *clpP* | 3 | Del-C | 365 | Gly122fs |
| *dnaK* | 3 | G-->A | 1115 | Ala372Val |
| *dacB* | 4 | C-->T | 343 | Gly115Ser |
| **CEFTAZIDIME/AVIBACTAM TREATMENT** |  |  |  |  |
| *dnaJ* | 5 | G-->A | 1081 | Pro361Ser |
| *pepA* | 6 | C-->T | 1439 | Gly480Asp |
| *ctpA* | 6 | G-->A | 971 | Ser324Asn |
| *glnD* | 6 | Ins-G | 2466-2467 | Asp823fs |
| *flgF* | 6 | C-->A | 676 | Glu226* |
| *pcm* | 8 | Ins-G | 530-531 | Arg180fs |
| *spoT* | 8 | T-->C | 931 | Phe311Leu |
| *PA14_45870* | 5 | Del-21 pb | 712-732 | Glu238fs |
| *PA14_45880* | 7 | C-->A | 159 | Met53Ile |
|  |  | G-->A | 1001 | Ser334Leu |
| *PA14_45890* | 7 | G-->A | 836 | Pro279Leu |
|  | 5 | G-->A | 830 | Ala277Val |
| INTERGENIC MUTATIONS |  |  |  |  |
| Between<br>*PA14_45410* and<br>*PA14_45430* | 10 | C-->T | 4044801 | - |
| Between 16S<br>ribosomal RNA<br>and *PA14_70920* | 12 | T-->C | 6317713 | - |
| Between *grpE* and<br>*recN* | 1 | T-->C | 5621761 | - |
| Between *mexR* and<br>*mexA* | 2 | C-->T | 486683 | - |

**Supplementary Table S2.**
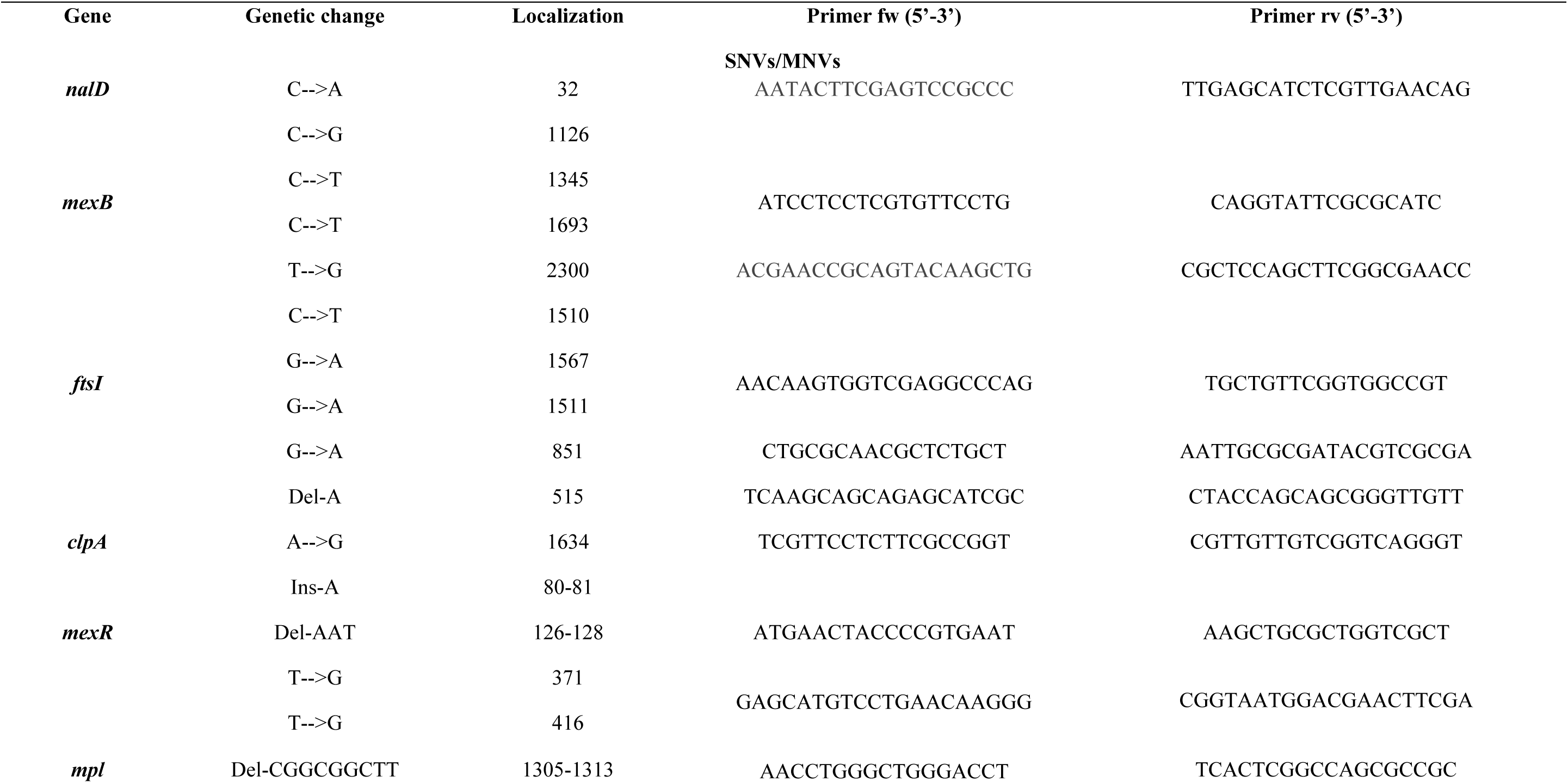
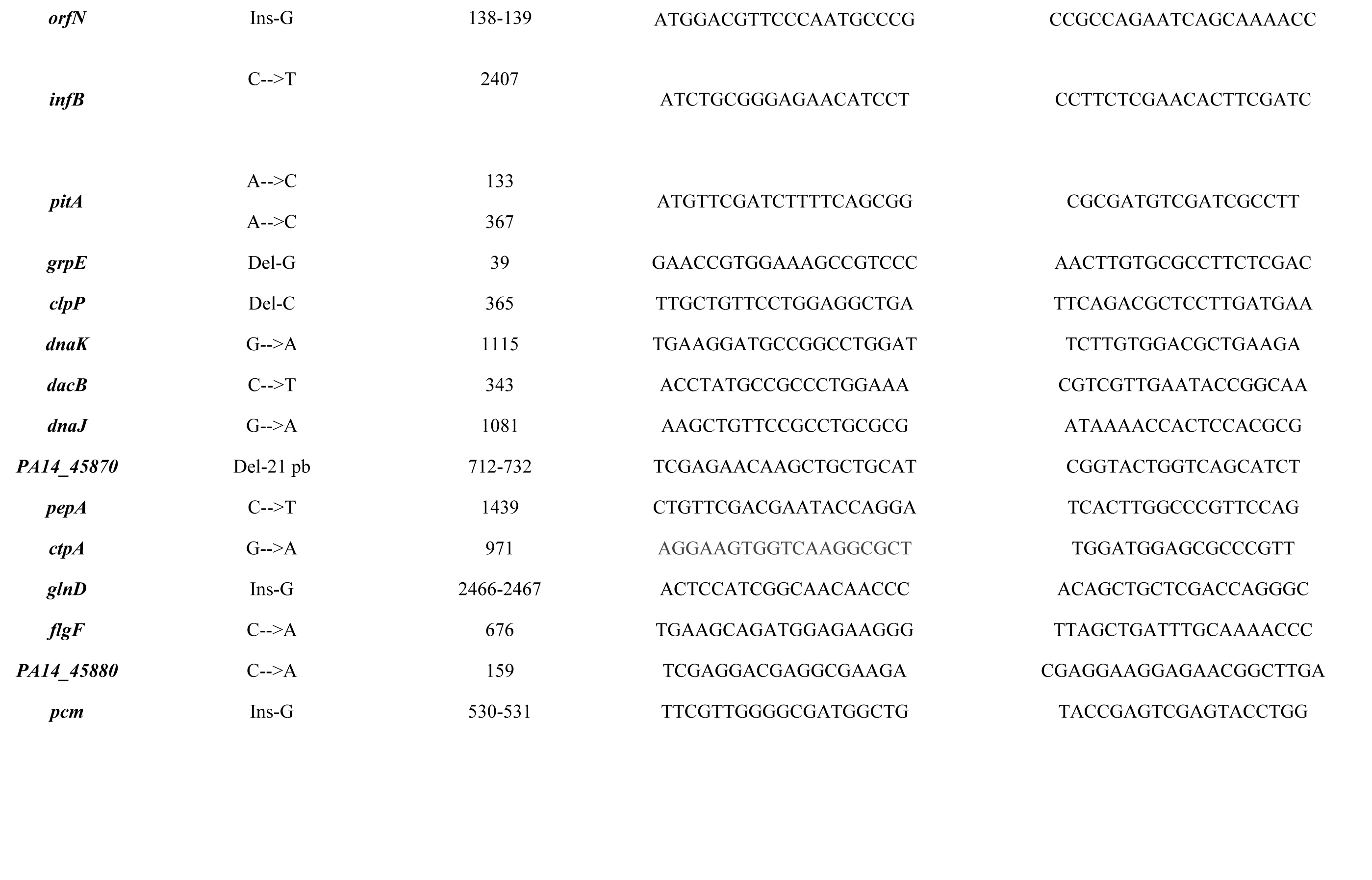
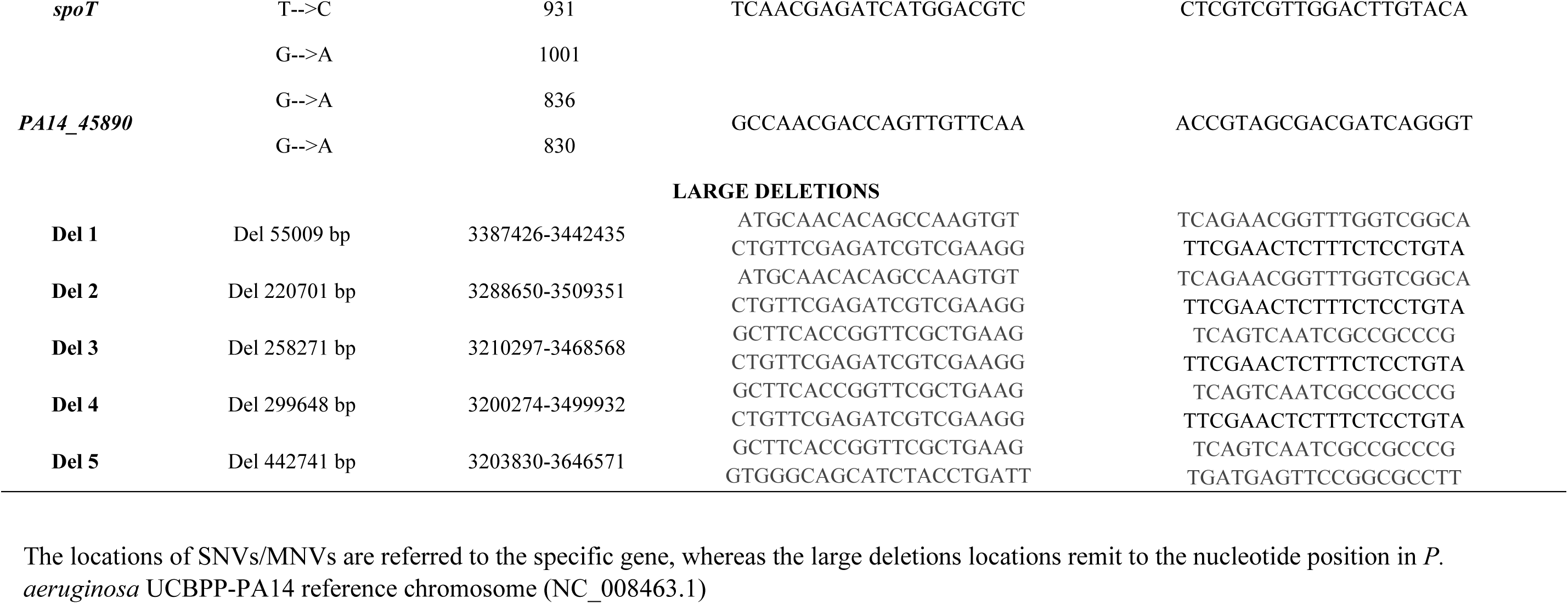
Primers used to verify nucleotide modifications detected in whole-genome sequencing. The locations of SNVs/MNVs are referred to the specific gene, whereas the large deletions locations remit to the nucleotide position in *P. aeruginosa* UCBPP-PA14 reference chromosome (NC_008463.1)

**Supplementary Table S3.** MIC values (µg/ml) obtained by double dilution for the population replicates during selective pressure with either ceftazidime or ceftazidime/avibactam. The table shows the liquid MIC values against ceftazidime for each replicate population every five days (the antibiotic concentration was doubled every 5 days). The grey boxes indicate the steps in which these particular replicates were unable to grow. In these situations, the previous ceftazidime concentrations (8MIC in populations 6, 8; and 16MIC in populations 3, 7) were maintained for the rest of the experiment.

| Treatment | Replicate | 5 d | 10 d | 15 d | 20 d | 25 d | 30 d |
| --- | --- | --- | --- | --- | --- | --- | --- |
| CEFTAZIDIME | 1 | 20 | 40 | 75 | 100 | 300 | 300 |
|  | 2 | 20 | 60 | 250 | 300 | 400 | 400 |
|  | 3 | 20 | 40 | 75 | 100 | 150 | 150 |
|  | 4 | 25 | 40 | 100 | 400 | 300 | 400 |
| CEFTAZIDIME<br>+<br>AVIBACTAM | 5 | 40 | 50 | 150 | 250 | 300 | 400 |
|  | 6 | 15 | 30 | 75 | 250 | 400 | 200 |
|  | 7 | 20 | 80 | 200 | 400 | 300 | 400 |
|  | 8 | 15 | 40 | 50 | 75 | 150 | 150 |
| CONTROLS | 9 | 4 | 4 | 4 | 4 | 4 | 4 |
|  | 10 | 4 | 4 | 4 | 4 | 4 | 4 |
|  | 11 | 4 | 4 | 4 | 4 | 4 | 4 |
|  | 12 | 4 | 4 | 4 | 4 | 4 | 4 |

**Supplementary Table S4.**
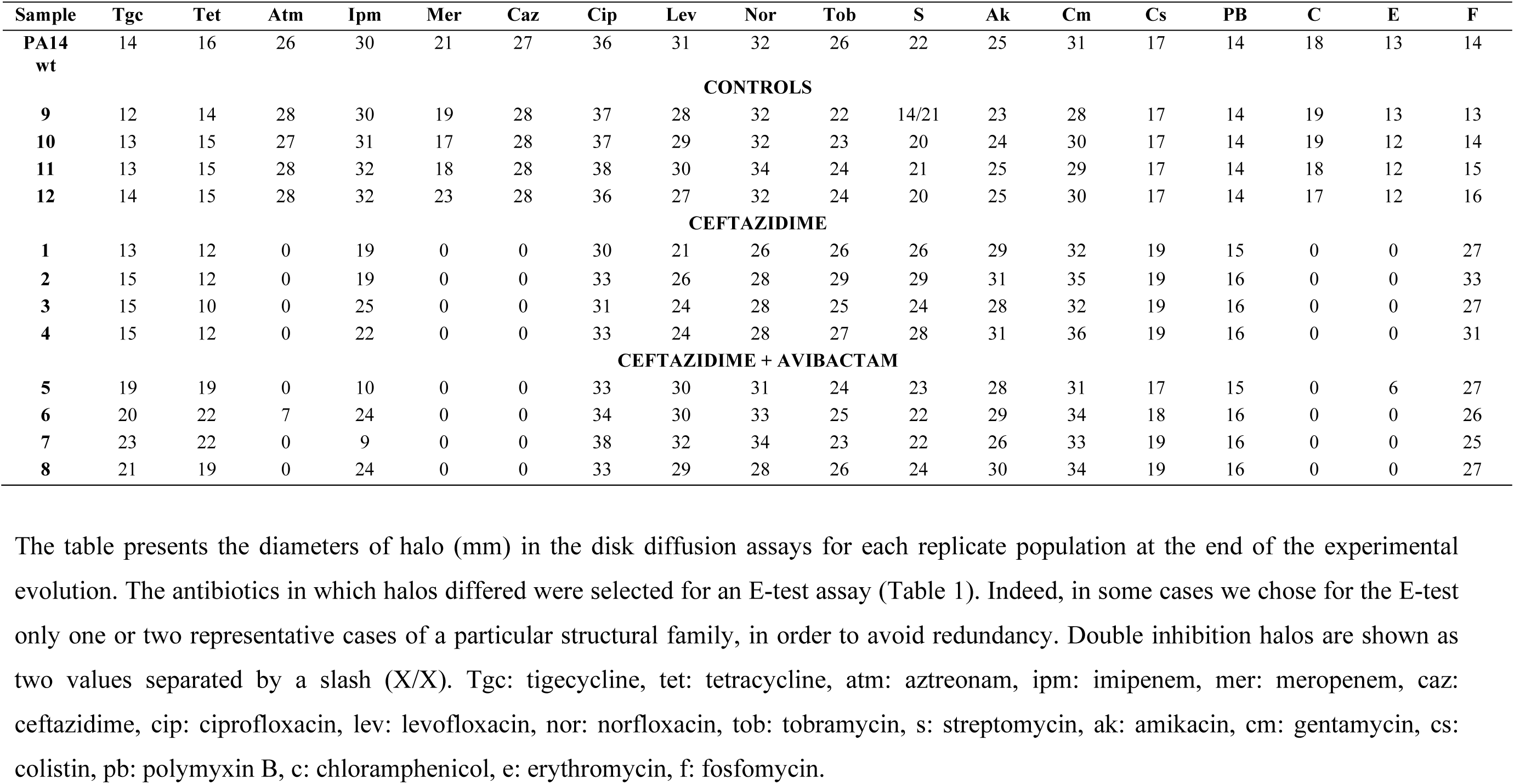
Disk diffusion tests with antibiotics of distinct structural families in the ceftazidime and ceftazidime/avibactam final evolved populations. The table presents the diameters of halo (mm) in the disk diffusion assays for each replicate population at the end of the experimental evolution. The antibiotics in which halos differed were selected for an E-test assay (Table 1). Indeed, in some cases we chose for the E-test only one or two representative cases of a particular structural family, in order to avoid redundancy. Double inhibition halos are shown as two values separated by a slash (X/X). Tgc: tigecycline, tet: tetracycline, atm: aztreonam, ipm: imipenem, mer: meropenem, caz: ceftazidime, cip: ciprofloxacin, lev: levofloxacin, nor: norfloxacin, tob: tobramycin, s: streptomycin, ak: amikacin, cm: gentamycin, cs: colistin, pb: polymyxin B, c: chloramphenicol, e: erythromycin, f: fosfomycin.

## REFERENCES

1. Silby MW, Winstanley C, Godfrey SA, Levy SB, Jackson RW. 2011. Pseudomonas genomes: diverse and adaptable. FEMS Microbiol Rev 35:652–80.

2. Martinez-Solano L, Macia MD, Fajardo A, Oliver A, Martinez JL. 2008. Chronic Pseudomonas aeruginosa infection in chronic obstructive pulmonary disease. Clinical Infectious Diseases 47:1526–33.

3. Talwalkar JS, Murray TS. 2016. The Approach to Pseudomonas aeruginosa in Cystic Fibrosis. Clin Chest Med 37:69–81.

4. Vila J, Martínez JL. 2008. Clinical impact of the over-expression of efflux pump in nonfermentative Gram-negative bacilli, development of efflux pump inhibitors. Current drug targets 9:797–807.

5. Oliver A, Mulet X, Lopez-Causape C, Juan C. 2015. The increasing threat of Pseudomonas aeruginosa high-risk clones. Drug Resist Updat 21-22:41–59.

6. Tzouvelekis LS, Markogiannakis A, Psichogiou M, Tassios PT, Daikos GL. 2012. Carbapenemases in Klebsiella pneumoniae and other Enterobacteriaceae: an evolving crisis of global dimensions. Clin Microbiol Rev 25:682–707.

7. Woodford N, Turton JF, Livermore DM. 2011. Multiresistant Gram-negative bacteria: the role of high-risk clones in the dissemination of antibiotic resistance. FEMS Microbiol Rev 35:736–55.

8. Buynak JD. 2006. Understanding the longevity of the beta-lactam antibiotics and of antibiotic/beta-lactamase inhibitor combinations. Biochem Pharmacol 71:930–40.

9. Liscio JL, Mahoney MV, Hirsch EB. 2015. Ceftolozane/tazobactam and ceftazidime/avibactam: two novel beta-lactam/beta-lactamase inhibitor combination agents for the treatment of resistant Gram-negative bacterial infections. Int J Antimicrob Agents 46:266–71.

10. Coleman K. 2011. Diazabicyclooctanes (DBOs): a potent new class of non-beta-lactam beta-lactamase inhibitors. Curr Opin Microbiol 14:550–5.

11. Aktas Z, Kayacan C, Oncul O. 2012. In vitro activity of avibactam (NXL104) in combination with beta-lactams against Gram-negative bacteria, including OXA-48 beta-lactamase-producing Klebsiella pneumoniae. Int J Antimicrob Agents 39:86–9.

12. Hayes MV, Orr DC. 1983. Mode of action of ceftazidime: affinity for the penicillin-binding proteins of Escherichia coli K12, Pseudomonas aeruginosa and Staphylococcus aureus. J Antimicrob Chemother 12:119–26.

13. Hidalgo JA, Vinluan CM, Antony N. 2016. Ceftazidime/avibactam: a novel cephalosporin/nonbeta-lactam beta-lactamase inhibitor for the treatment of complicated urinary tract infections and complicated intra-abdominal infections. Drug Des Devel Ther 10:2379–86.

14. Pitart C, Marco F, Keating TA, Nichols WW, Vila J. 2015. Activity of ceftazidime-avibactam against fluoroquinolone-resistant Enterobacteriaceae and Pseudomonas aeruginosa. Antimicrob Agents Chemother 59:3059–65.

15. Calvopina K, Hinchliffe P, Brem J, Heesom KJ, Johnson S, Cain R, Lohans CT, Fishwick CWG, Schofield CJ, Spencer J, Avison MB. 2017. Structural/mechanistic insights into the efficacy of nonclassical beta-lactamase inhibitors against extensively drug resistant Stenotrophomonas maltophilia clinical isolates. Mol Microbiol 106:492–504.

16. Lopez-Hernandez I, Alonso N, Fernandez-Martinez M, Zamorano L, Rivera A, Oliver A, Conejo MC, Martinez-Martinez L, Navarro F, Pascual A. 2017. Activity of ceftazidime-avibactam against multidrug-resistance Enterobacteriaceae expressing combined mechanisms of resistance. Enferm Infecc Microbiol Clin 35:499–504.

17. Fraile-Ribot PA, Cabot G, Mulet X, Perianez L, Martin-Pena ML, Juan C, Perez JL, Oliver A. 2017. Mechanisms leading to in vivo ceftolozane/tazobactam resistance development during the treatment of infections caused by MDR Pseudomonas aeruginosa. J Antimicrob Chemother doi:10.1093/jac/dkx424.

18. Zhanel GG, Lawson CD, Adam H, Schweizer F, Zelenitsky S, Lagace-Wiens PR, Denisuik A, Rubinstein E, Gin AS, Hoban DJ, Lynch JP, 3rd, Karlowsky JA. 2013. Ceftazidime-avibactam: a novel cephalosporin/beta-lactamase inhibitor combination. Drugs 73:159–77.

19. Levin BR, Rozen DE. 2006. Non-inherited antibiotic resistance. Nat Rev Microbiol 4:556–62.

20. Martinez JL, Rojo F. 2011. Metabolic regulation of antibiotic resistance. FEMS Microbiol Rev 35:768–89.

21. Martinez JL, Fajardo A, Garmendia L, Hernandez A, Linares JF, Martinez-Solano L, Sanchez MB. 2009. A global view of antibiotic resistance. FEMS Microbiol Rev 33:44–65.

22. Rodriguez-Rojas A, Mena A, Martin S, Borrell N, Oliver A, Blazquez J. 2009. Inactivation of the hmgA gene of Pseudomonas aeruginosa leads to pyomelanin hyperproduction, stress resistance and increased persistence in chronic lung infection. Microbiology 155:1050–7.

23. Alvarez-Ortega C, Wiegand I, Olivares J, Hancock RE, Martinez JL. 2010. Genetic determinants involved in the susceptibility of Pseudomonas aeruginosa to beta-lactam antibiotics. Antimicrob Agents Chemother 54:4159–67.

24. Masuda N, Sakagawa E, Ohya S, Gotoh N, Tsujimoto H, Nishino T. 2000. Substrate specificities of MexAB-OprM, MexCD-OprJ, and MexXY-oprM efflux pumps in Pseudomonas aeruginosa. Antimicrob Agents Chemother 44:3322–7.

25. Morita Y, Cao L, Gould VC, Avison MB, Poole K. 2006.nalD encodes a second repressor of the mexAB-oprM multidrug efflux operon of Pseudomonas aeruginosa. J Bacteriol 188:8649–54.

26. Sobel ML, Hocquet D, Cao L, Plesiat P, Poole K. 2005. Mutations in PA3574 (nalD) lead to increased MexAB-OprM expression and multidrug resistance in laboratory and clinical isolates of Pseudomonas aeruginosa. Antimicrob Agents Chemother 49:1782–6.

27. Cabot G, Ocampo-Sosa AA, Dominguez MA, Gago JF, Juan C, Tubau F, Rodriguez C, Moya B, Pena C, Martinez-Martinez L, Oliver A, Spanish Network for Research in Infectious D. 2012. Genetic markers of widespread extensively drug-resistant Pseudomonas aeruginosa high-risk clones. Antimicrob Agents Chemother 56:6349–57.

28. Jorth P, McLean K, Ratjen A, Secor PR, Bautista GE, Ravishankar S, Rezayat A, Garudathri J, Harrison JJ, Harwood RA, Penewit K, Waalkes A, Singh PK, Salipante SJ. 2017. Evolved Aztreonam Resistance Is Multifactorial and Can Produce Hypervirulence in Pseudomonas aeruginosa. MBio 8.

29. Cabot G, Lopez-Causape C, Ocampo-Sosa AA, Sommer LM, Dominguez MA, Zamorano L, Juan C, Tubau F, Rodriguez C, Moya B, Pena C, Martinez-Martinez L, Plesiat P, Oliver A. 2016. Deciphering the Resistome of the Widespread Pseudomonas aeruginosa Sequence Type 175 International High-Risk Clone through Whole-Genome Sequencing. Antimicrob Agents Chemother 60:7415–7423.

30. Lopez-Causape C, Sommer LM, Cabot G, Rubio R, Ocampo-Sosa AA, Johansen HK, Figuerola J, Canton R, Kidd TJ, Molin S, Oliver A. 2017. Evolution of the Pseudomonas aeruginosa mutational resistome in an international Cystic Fibrosis clone. Sci Rep 7:5555.

31. Kos VN, Deraspe M, McLaughlin RE, Whiteaker JD, Roy PH, Alm RA, Corbeil J, Gardner H. 2015. The resistome of Pseudomonas aeruginosa in relationship to phenotypic susceptibility. Antimicrob Agents Chemother 59:427–36.

32. Fernandez L, Breidenstein EB, Song D, Hancock RE. 2012. Role of intracellular proteases in the antibiotic resistance, motility, and biofilm formation of Pseudomonas aeruginosa. Antimicrob Agents Chemother 56:1128–32.

33. Aguilera Rossi CG, Gomez-Puertas P, Ayala Serrano JA. 2016. In vivo functional and molecular characterization of the Penicillin-Binding Protein 4 (DacB) of Pseudomonas aeruginosa. BMC Microbiol 16:234.

34. Schirm M, Arora SK, Verma A, Vinogradov E, Thibault P, Ramphal R, Logan SM. 2004. Structural and genetic characterization of glycosylation of type a flagellin in Pseudomonas aeruginosa. J Bacteriol 186:2523–31.

35. Wong A, Rodrigue N, Kassen R. 2012. Genomics of adaptation during experimental evolution of the opportunistic pathogen Pseudomonas aeruginosa. PLoS Genet 8:e1002928.

36. Liu A, Tran L, Becket E, Lee K, Chinn L, Park E, Tran K, Miller JH. 2010. Antibiotic sensitivity profiles determined with an Escherichia coli gene knockout collection: generating an antibiotic bar code. Antimicrob Agents Chemother 54:1393–403.

37. Fajardo A, Martinez-Martin N, Mercadillo M, Galan JC, Ghysels B, Matthijs S, Cornelis P, Wiehlmann L, Tummler B, Baquero F, Martinez JL. 2008. The neglected intrinsic resistome of bacterial pathogens. PLoS ONE 3:e1619.

38. Kohler T, Michea-Hamzehpour M, Epp SF, Pechere JC. 1999. Carbapenem activities against Pseudomonas aeruginosa: respective contributions of OprD and efflux systems. Antimicrob Agents Chemother 43:424–7.

39. Hauser AR, Kang PJ, Engel JN. 1998. PepA, a secreted protein of Pseudomonas aeruginosa, is necessary for cytotoxicity and virulence. Mol Microbiol 27:807–18.

40. Potvin E, Lehoux DE, Kukavica-Ibrulj I, Richard KL, Sanschagrin F, Lau GW, Levesque RC. 2003. In vivo functional genomics of Pseudomonas aeruginosa for high-throughput screening of new virulence factors and antibacterial targets. Environ Microbiol 5:1294–308.

41. Isabella VM, Campbell AJ, Manchester J, Sylvester M, Nayar AS, Ferguson KE, Tommasi R, Miller AA. 2015. Toward the rational design of carbapenem uptake in Pseudomonas aeruginosa. Chem Biol 22:535–547.

42. Yen P, Papin JA. 2017. History of antibiotic adaptation influences microbial evolutionary dynamics during subsequent treatment. PLoS Biol 15:e2001586.

43. Homma M, Kutsukake K, Hasebe M, Iino T, Macnab RM. 1990. FlgB, FlgC, FlgF and FlgG. A family of structurally related proteins in the flagellar basal body of Salmonella typhimurium. J Mol Biol 211:465–77.

44. Seo J, Darwin AJ. 2013. The Pseudomonas aeruginosa periplasmic protease CtpA can affect systems that impact its ability to mount both acute and chronic infections. Infect Immun 81:4561–70.

45. Contreras A, Drummond M, Bali A, Blanco G, Garcia E, Bush G, Kennedy C, Merrick M. 1991. The product of the nitrogen fixation regulatory gene nfrX of Azotobacter vinelandii is functionally and structurally homologous to the uridylyltransferase encoded by glnD in enteric bacteria. J Bacteriol 173:7741–9.

46. Reading C, Cole M. 1977. Clavulanic acid: a beta-lactamase-inhiting beta-lactam from Streptomyces clavuligerus. Antimicrob Agents Chemother 11:852–7.

47. Baquero F, Reig M. 1989. Mechanisms of antimicrobial resistance in anaerobic bacteria: the predictive approach. Scand J Infect Dis Suppl 62:25–8.

48. Blazquez J, Baquero MR, Canton R, Alos I, Baquero F. 1993. Characterization of a new TEM-type beta-lactamase resistant to clavulanate, sulbactam, and tazobactam in a clinical isolate of Escherichia coli. Antimicrob Agents Chemother 37:2059–63.

49. Canton R, Gonzalez-Alba JM, Galan JC. 2012. CTX-M Enzymes: Origin and Diffusion. Front Microbiol 3:110.

50. Toussaint KA, Gallagher JC. 2015. beta-lactam/beta-lactamase inhibitor combinations: from then to now. Ann Pharmacother 49:86–98.

51. Cabot G, Zamorano L, Moya B, Juan C, Navas A, Blazquez J, Oliver A. 2016. Evolution of Pseudomonas aeruginosa Antimicrobial Resistance and Fitness under Low and High Mutation Rates. Antimicrob Agents Chemother 60:1767–78.

52. Mayer-Hamblett N, Rosenfeld M, Gibson RL, Ramsey BW, Kulasekara HD, Retsch-Bogart GZ, Morgan W, Wolter DJ, Pope CE, Houston LS, Kulasekara BR, Khan U, Burns JL, Miller SI, Hoffman LR. 2014. Pseudomonas aeruginosa in vitro phenotypes distinguish cystic fibrosis infection stages and outcomes. Am J Respir Crit Care Med 190:289–97.

53. Shen M, Zhang H, Shen W, Zou Z, Lu S, Li G, He X, Agnello M, Shi W, Hu F, Le S. 2018. Pseudomonas aeruginosa MutL promotes large chromosomal deletions through non-homologous end joining to prevent bacteriophage predation. Nucleic Acids Research 46:4505–4514.

54. Hocquet D, Petitjean M, Rohmer L, Valot B, Kulasekara HD, Bedel E, Bertrand X, Plesiat P, Kohler T, Pantel A, Jacobs MA, Hoffman LR, Miller SI. 2016. Pyomelanin-producing Pseudomonas aeruginosa selected during chronic infections have a large chromosomal deletion which confers resistance to pyocins. Environ Microbiol 18:3482–3493.

55. Chalhoub H, Saenz Y, Rodriguez-Villalobos H, Denis O, Kahl BC, Tulkens PM, Van Bambeke F, Pan YP, Xu YH, Wang ZX, Fang YP, Shen JL, Riou M, Avrain L, Carbonnelle S, El Garch F, Pirnay JP, De Vos D, Plesiat P, Tulkens PM, Van Bambeke F, Castanheira M, Mills JC, Farrell DJ, Jones RN. High-level resistance to meropenem in clinical isolates of Pseudomonas aeruginosa in the absence of carbapenemases: role of active efflux and porin alterations Overexpression of MexAB-OprM efflux pump in carbapenem-resistant Pseudomonas aeruginosa Increase of efflux-mediated resistance in Pseudomonas aeruginosa during antibiotic treatment in patients suffering from nosocomial pneumonia Mutation-driven beta-lactam resistance mechanisms among contemporary ceftazidime-nonsusceptible Pseudomonas aeruginosa isolates from U.S. hospitals.

56. Pan YP, Xu YH, Wang ZX, Fang YP, Shen JL. Overexpression of MexAB-OprM efflux pump in carbapenem-resistant Pseudomonas aeruginosa.

57. Riou M, Avrain L, Carbonnelle S, El Garch F, Pirnay JP, De Vos D, Plesiat P, Tulkens PM, Van Bambeke F. Increase of efflux-mediated resistance in Pseudomonas aeruginosa during antibiotic treatment in patients suffering from nosocomial pneumonia.

58. Castanheira M, Mills JC, Farrell DJ, Jones RN. Mutation-driven beta-lactam resistance mechanisms among contemporary ceftazidime-nonsusceptible Pseudomonas aeruginosa isolates from U.S. hospitals.

59. Mulet X, Moya B, Juan C, Macia MD, Perez JL, Blazquez J, Oliver A. 2011. Antagonistic interactions of Pseudomonas aeruginosa antibiotic resistance mechanisms in planktonic but not biofilm growth. Antimicrob Agents Chemother 55:4560–8.

60. Lahiri SD, Walkup GK, Whiteaker JD, Palmer T, McCormack K, Tanudra MA, Nash TJ, Thresher J, Johnstone MR, Hajec L, Livchak S, McLaughlin RE, Alm RA. 2015. Selection and molecular characterization of ceftazidime/avibactam-resistant mutants in Pseudomonas aeruginosa strains containing derepressed AmpC. J Antimicrob Chemother 70:1650–8.

61. Blair JM, Bavro VN, Ricci V, Modi N, Cacciotto P, Kleinekathfer U, Ruggerone P, Vargiu AV, Baylay AJ, Smith HE, Brandon Y, Galloway D, Piddock LJ. 2015. AcrB drug-binding pocket substitution confers clinically relevant resistance and altered substrate specificity. Proc Natl Acad Sci U S A 112:3511–6.

62. Rau MH, Marvig RL, Ehrlich GD, Molin S, Jelsbak L. 2012. Deletion and acquisition of genomic content during early stage adaptation of Pseudomonas aeruginosa to a human host environment. Environ Microbiol 14:2200–11.

63. Alvarez-Ortega C, Wiegand I, Olivares J, Hancock RE, Martinez JL. 2011. The intrinsic resistome of Pseudomonas aeruginosa to beta-lactams. Virulence 2:144–6.

